## Supplementary material for "Matryoshka RNA virus 1: a novel RNA virus associated with *Plasmodium* parasites in human malaria"

### Supplementary Figures and Tables

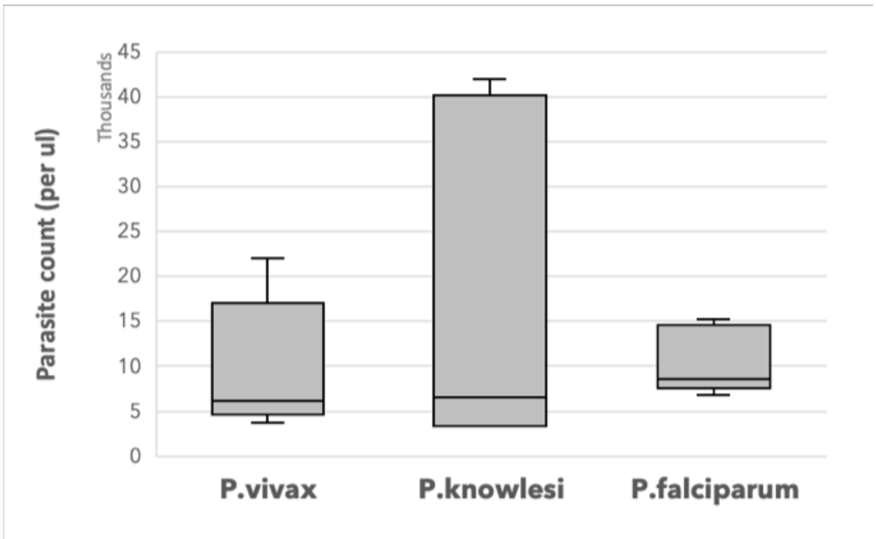


**Figure S1.** ***Plasmodium*** **parasite count in human blood samples.** Parasite counts are expressed as the number of parasites per µl of blood.

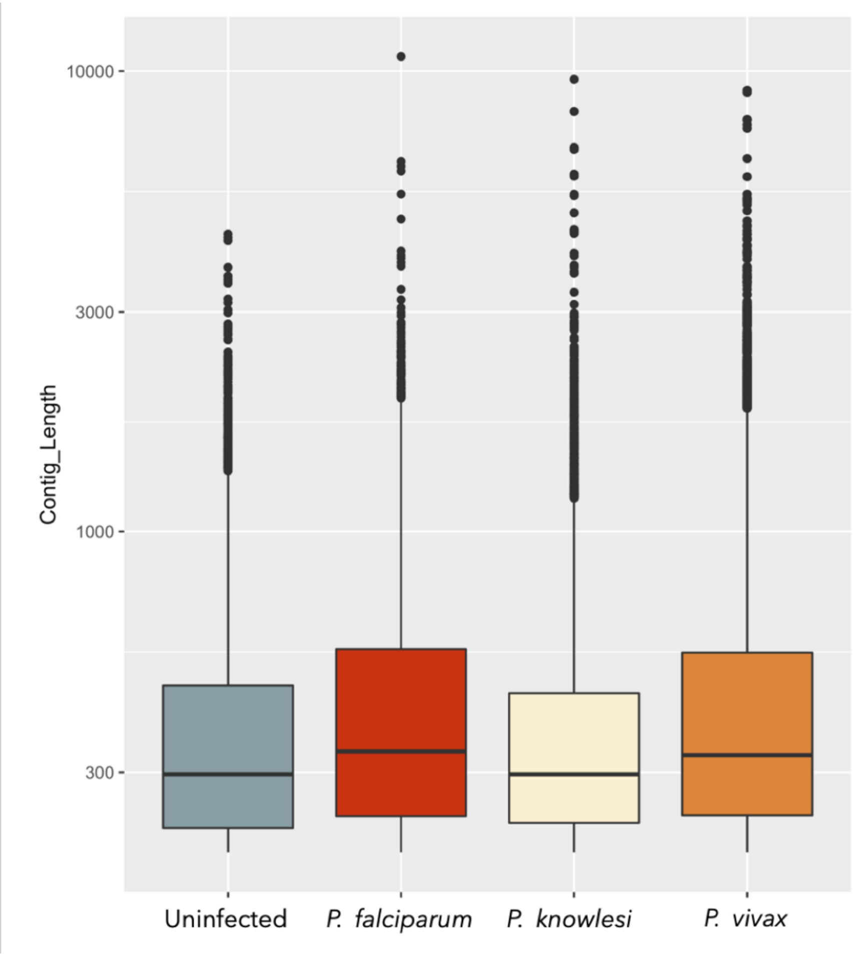


**Figure S2. Trinity assembly results.** Contig length and count obtained after performing Trinity assembly of libraries depleted in rRNA, human and *Plasmodium* reads.

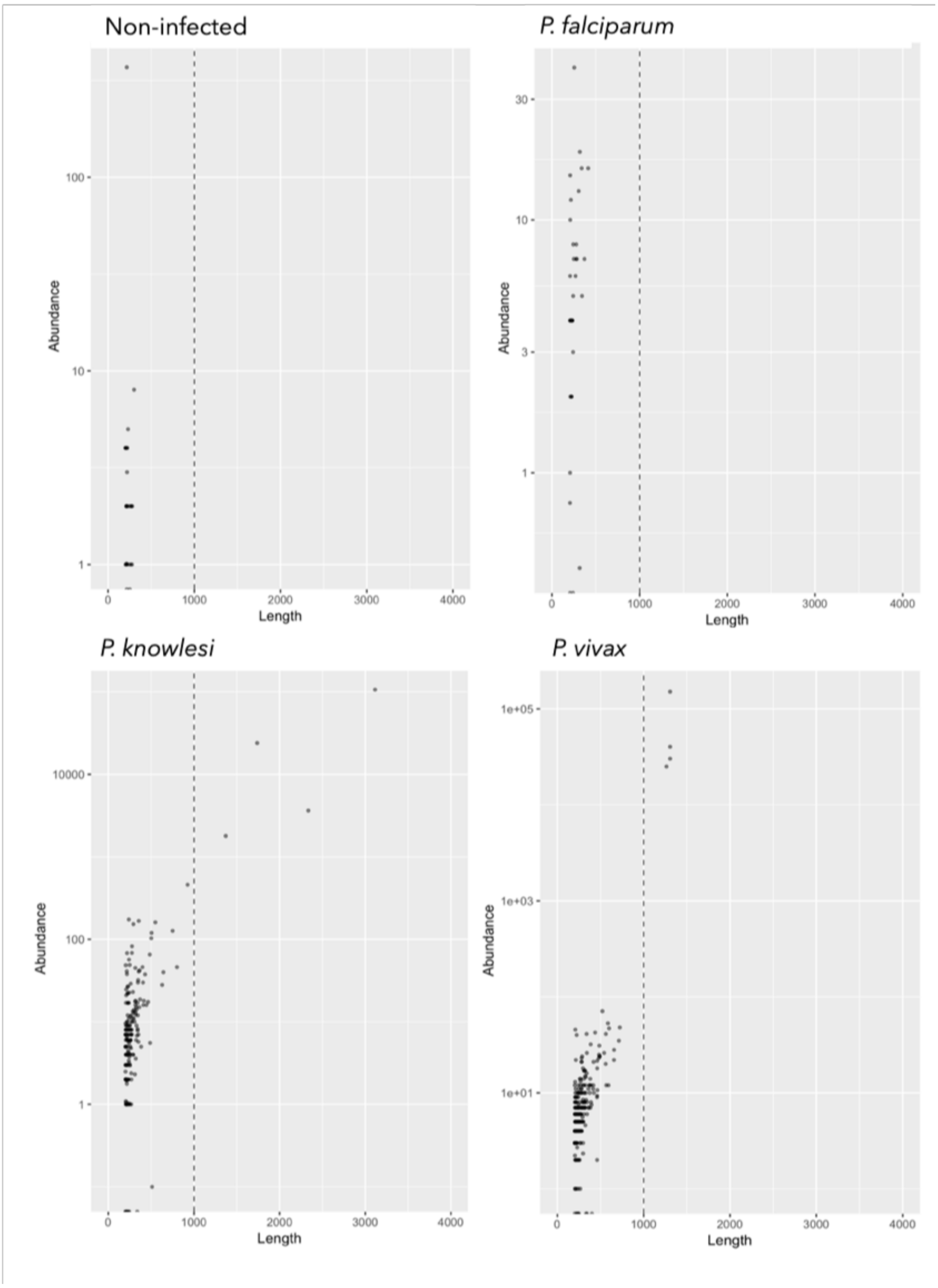


**Figure S3. Orphan contig length and abundance.** A 1000nt cut-off is applied to identify candidate RNA viruses. Abundance is expressed using the expected count value provided by the RSEM analysis.

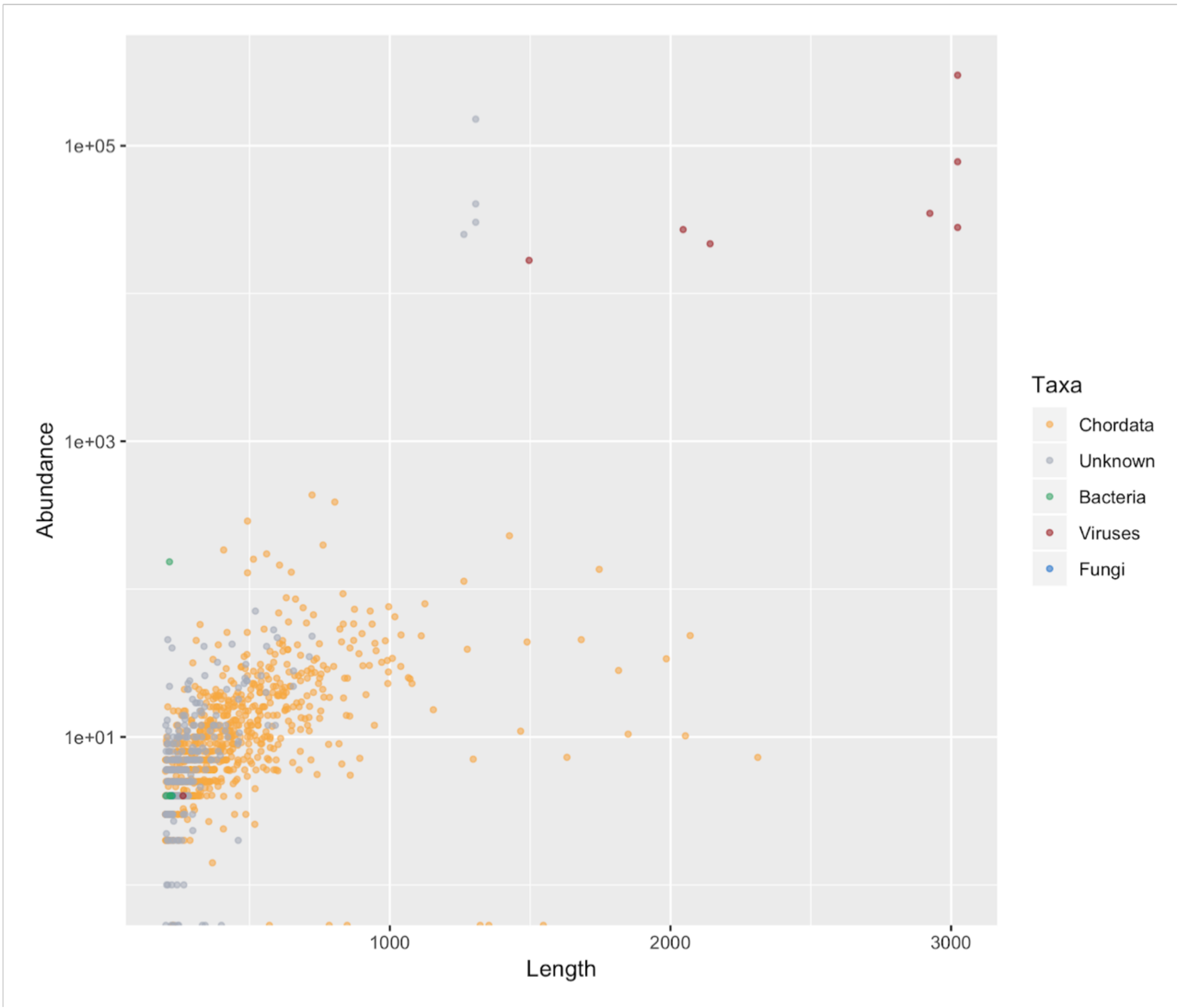


**Figure S4. Length and abundance of non-major host candidate contigs (Blastn/Blastx).** Abundance is expressed using the expected count value provided by the RSEM analysis.

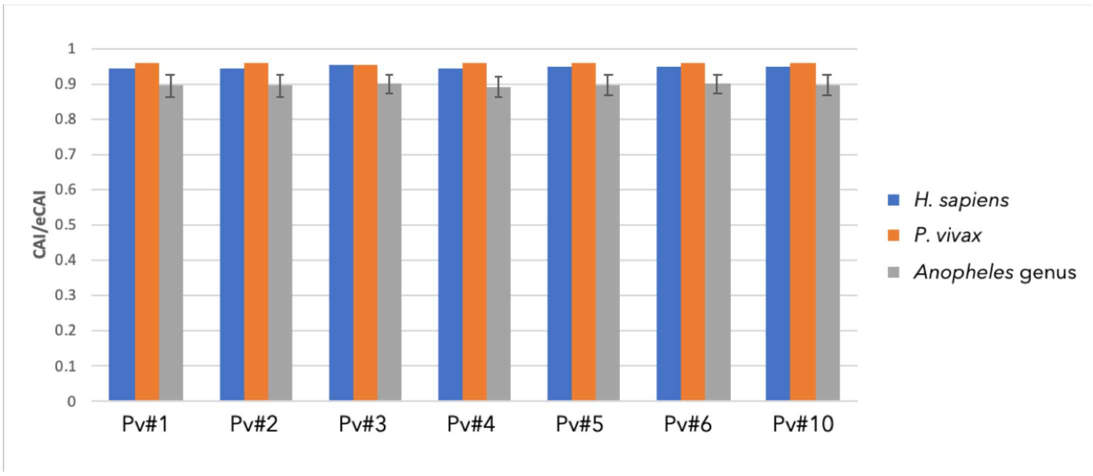


**Figure S5.** Comparison of CAI values obtained by comparing codon usage of MaRNAV-1 viral contigs retrieved from each *Plasmodium* library to the potential hosts *P. vivax*, *H. sapiens* and mosquitoes of the genus *Anopheles*.

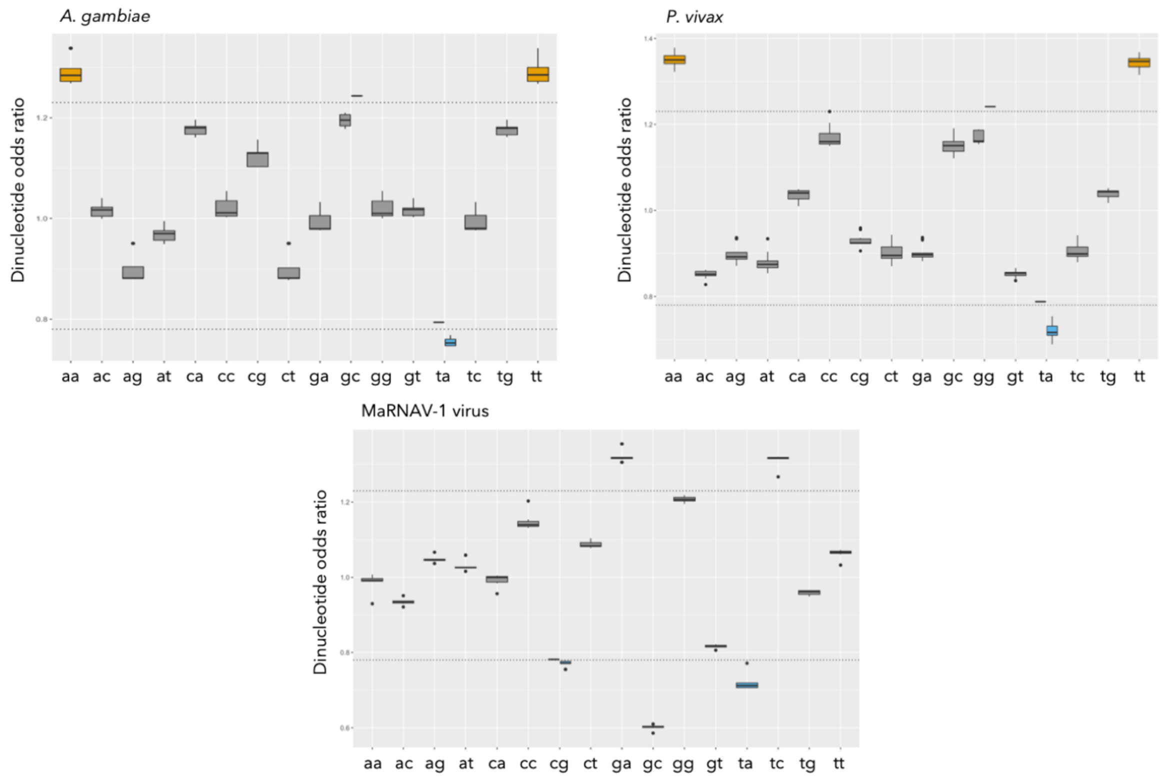


**Figure S6.** Odds ratios of dinucleotides (fxy/fxfy) obtained from MaRNAV-1 contigs (bottom) versus *P. vivax* (top, right) and *A. gambiae* (top, left) genomic sequence.

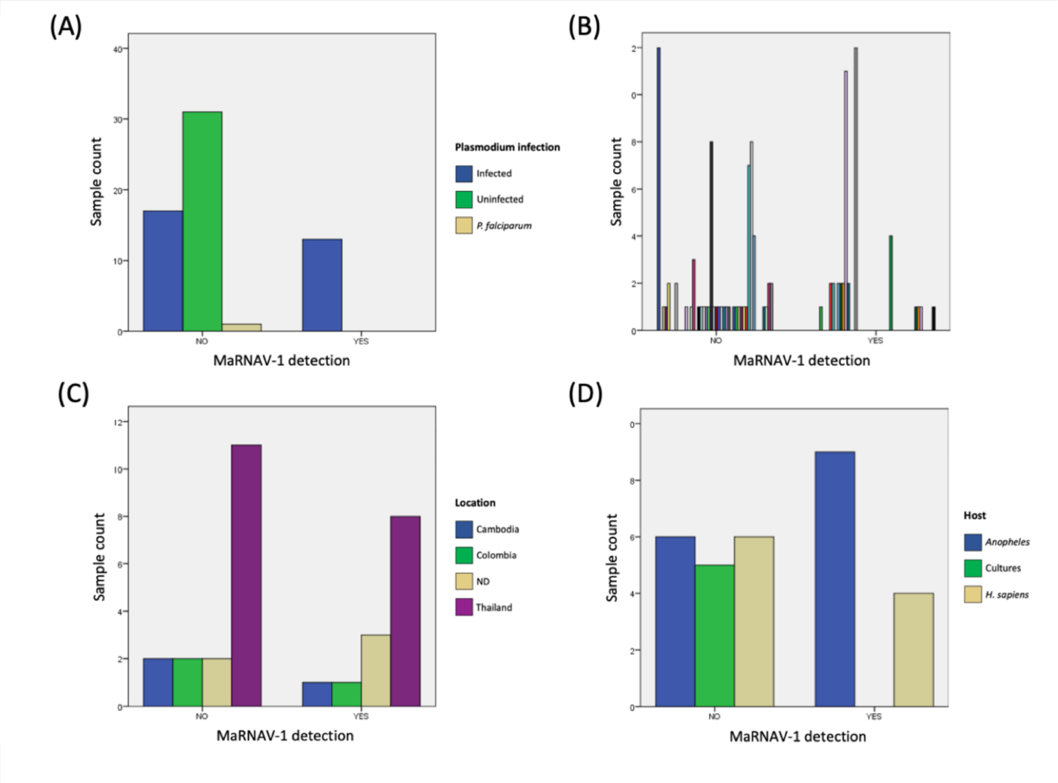


**Figure S7.** Association study between MaRNAV-1 contigs and the characteristics of *P. vivax* libraries at the SRA (Chi-squared tests). (A) *Plasmodium* infection association test. (B) Biological replicates association test. Replicates corresponding to the same biological sample are grouped by color. (C) Sample location association test. (D) Host association test.

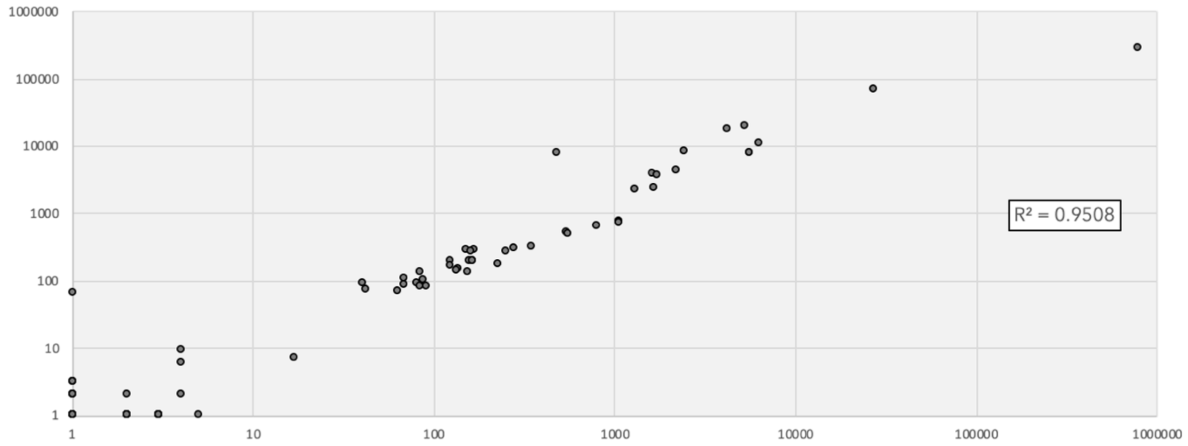


**Figure S8.** Read count mapping to the MaRNAV-1 segment I (x-axis, log scaled) and Segment II (y-axis, log scaled) in *P. vivax* SRA data sets. The R-squared value is indicated on the right.

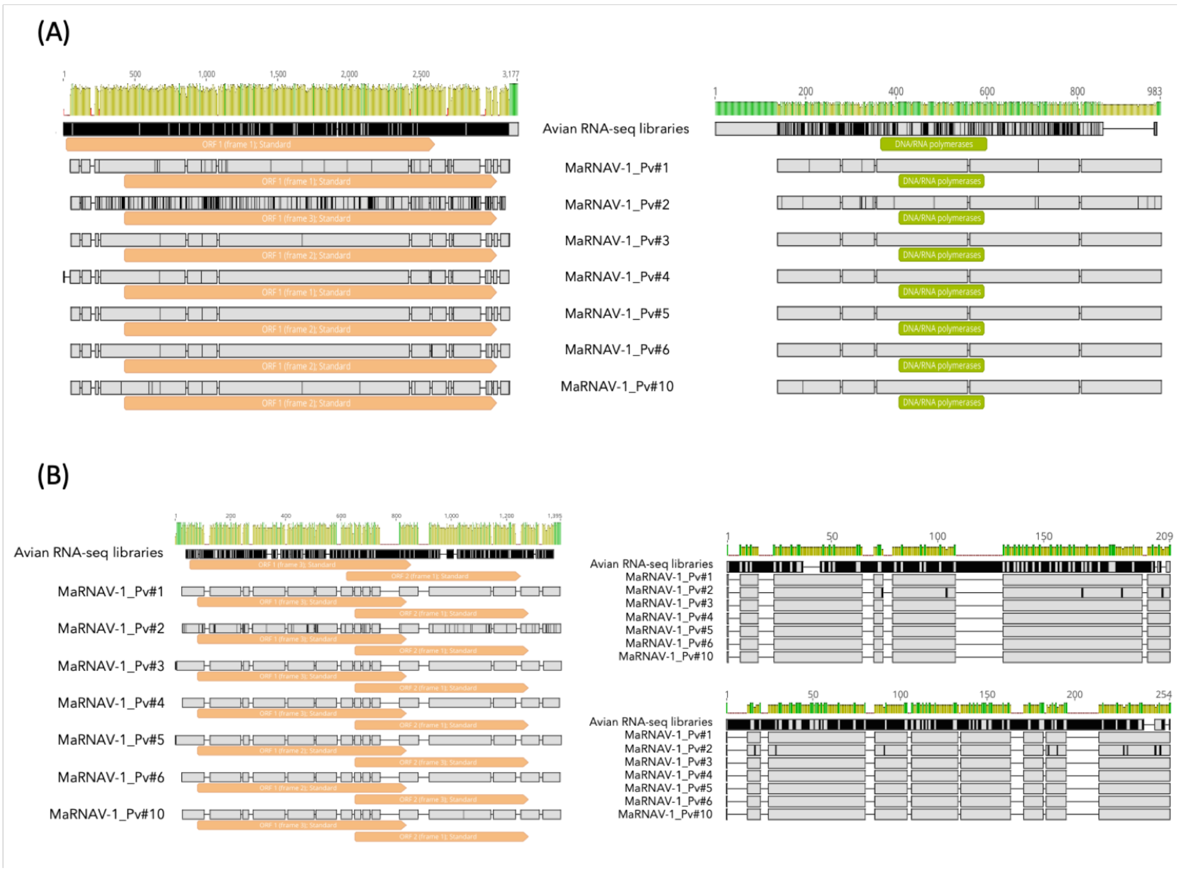


**Figure S9. Sequence alignment (MAFFT) of MaRNAV-2 contigs in *Leucocytozoon*-infected bird and MaRNAV-1 sequences in *P. vivax*-infected humans.** (A) Analysis of segment I. (B) Analysis of segment II. Nucleotide alignments are shown on the left and protein alignments are shown on the right (top - ORF1; bottom - ORF2). Orange boxes: predicted ORF using standard genetic code. Light green boxes: InterProScan domain prediction. Yellow to green plots: level of sequence conservation.

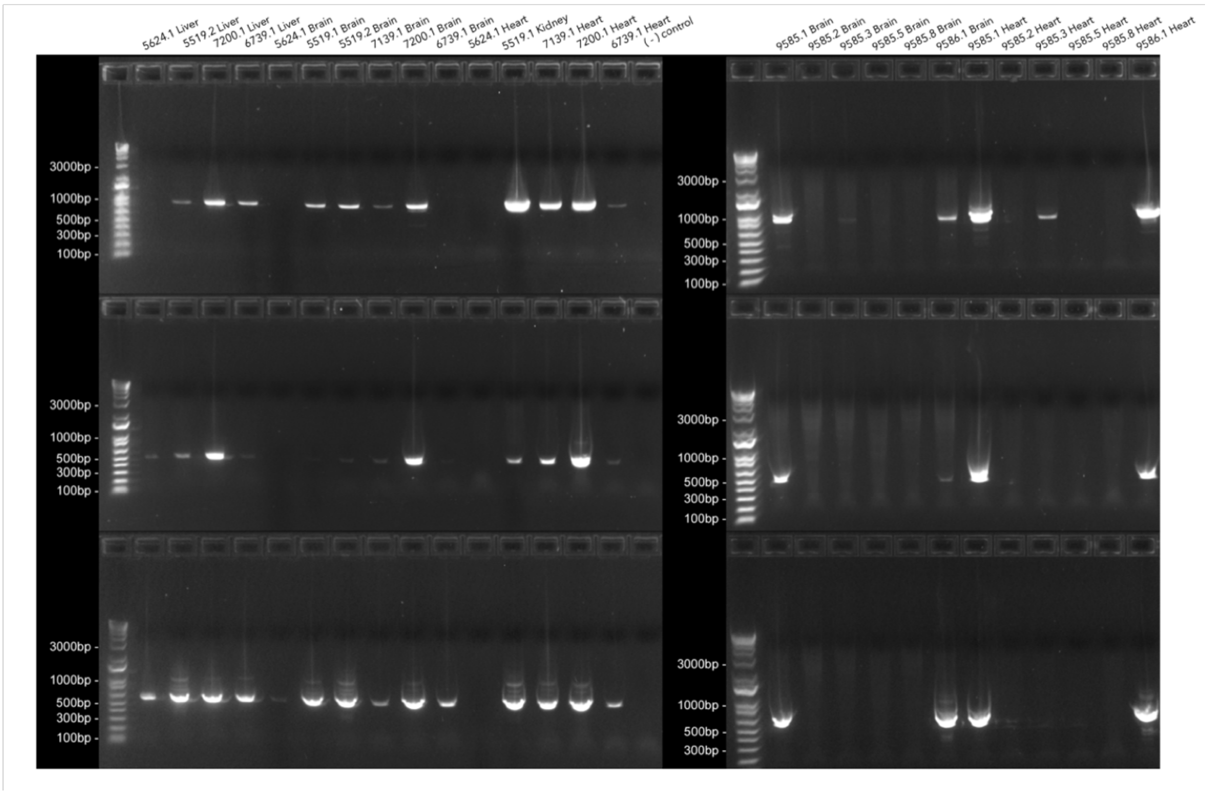


**Figure S10.** PCR-based detection of leucocytozoons and MaRNAV-2 (2 segments) from avian cDNA samples. Top: *Leucocytozoon* CytB PCR; Middle: MaRNAV-2 segment I homolog detection; Bottom: MaRNAV-2 segment II homolog detection.

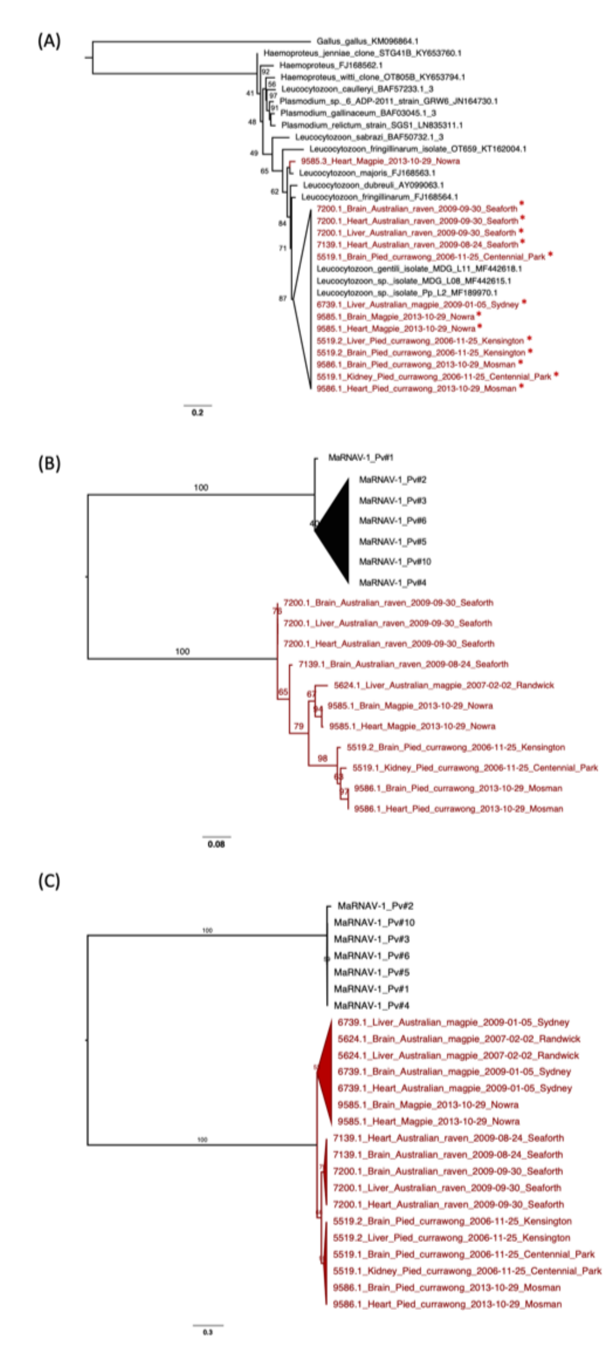


**Figure S11.** ML phylogenetic trees of parasites and MaRNAV-2 homologs obtained from bird samples. (A) Hematozoa CytB phylogeny. The CytB sequence from *Gallus gallus* is used as an outgroup. Samples positive for MaRNAV-2 I are marked with *. (B) MaRNAV-2 segment I phylogeny; (C) MaRNAV-2 segment II phylogeny. Sequences from bird samples are shown in red.

**Table S1. Description of human blood samples used in this study.** PCR: PCR-based validation of *Plasmodium* species using species-specific primers: Pv - *P. vivax*; Pk - *P. knowlesi*; Pf - *P. falciparum*; pc - parasite counting (i.e. parasite density / µL = number of parasites counted x patient's lab leukocyte result / 200 leukocytes counted).

| Sample | Study | Date | Village | District | PCR | pc |
|---|---|---|---|---|---|---|
| **1** | QDM015 | Apr-14 | Talas | Kota Marudu | Pv | 4551 |
| **2** | MV001 | Jan-13 | Bambangan Ulu | Kota Marudu | Pv | 16968 |
| **3** | MV002 | Jan-13 | Sonsogon Magandai | Kota Marudu | Pv | 21952 |
| **4** | MV039 | Nov-14 | Sinar 9 Bombong | Kota Marudu | Pv | 3694 |
| **5** | MV036 | Oct-14 | Kalibambang | Kota Marudu | Pv | 10590 |
| **6** | MV038 | Nov-14 | Sinar 9 Bombong | Kota Marudu | Pv | 6027 |
| **10** | QDM014 | Apr-14 | Sonsogon Magandai | Kota Marudu | Pv | 4950 |
| **18** | QDM027 | May-14 | Baru Malalin | Belarun | Pk | 39565 |
| **19** | MK011 | Apr-13 | Togudon | Kota Belud | Pk | 3245 |
| **20** | MK072 | Aug-14 | Malangkap | Kota Marudu | Pk | 7177 |
| **21** | KK042 | Apr-13 | Balak-Balak | Kudat | Pk | 3363 |
| **22** | KK077 | Jan-14 | Kusilad Darat | Kudat | Pk | 5674 |
| **27** | QDM032 | May-14 | Hatob | Kota Marudu | Pk | 41882 |
| **28** | QEM782 | Jan-14 | Damaran | Kudat | Pf | 8199 |
| **31** | MF024 | Sep-14 | Matanggal | Kota Marudu | Pf | 8514 |
| **32** | MF028 | Oct-14 | Salimandut | Kota Marudu | Pf | 15207 |
| **33** | MF029 | Nov-14 | Gana | Kota Marudu | Pf | 6783 |
| **35** | QDM056 | Sep-14 | Sonsogon Magandai | Kota Marudu | Pf | 13977 |
| **38** | QECK037 | Oct-14 | Pantai Bahagia | Kudat | Negative | - |
| **39** | QECK039 | Oct-14 | Perpaduan | Kudat | Negative | - |
| **40** | QECK042 | Oct-14 | Lok Tohog | Pitas | Negative | - |
| **42** | QECK038 | Oct-14 | Batu 1/2 Jalan Atas | Kudat | Negative | - |
| **45** | QECK040 | Oct-14 | Penaitan | Kudat | Negative | - |

| | | | | | | |
|---|---|---|---|---|---|---|
| **46** | QECK004 | Apr-13 | Taman Orkid Kudat | Kudat | Negative | - |

**Table S2. BioProject and corresponding SRA accessions used in this study.**

| Species | BioProject | No. libraries |
|---|---|---|
| ***P. chabaudi*** | PRJEB1284 | 60 |
| | PRJEB1500 | 24 |
| | PRJEB4572 | 84 |
| | PRJEB4731 | 102 |
| | PRJEB6241 | 78 |
| | PRJEB6242 | 22 |
| | **All** | **370** |
| ***P. cynomolgi*** | PRJNA356968 | 52 |
| | **All** | **52** |
| ***P. falciparum*** | PRJEB19245 | 441 |
| | PRJEB19644 | 60 |
| | PRJEB21707 | 60 |
| | PRJEB24218 | 128 |
| | PRJEB25413 | 9 |
| | PRJNA167166 | 3 |
| | PRJNA308455 | 3 |
| | PRJNA310220 | 12 |
| | PRJNA315899 | 6 |
| | PRJNA391508 | 16 |
| | PRJNA401189 | 4 |
| | **All** | **742** |
| ***P. knowlesi*** | PRJEB24220 | 69 |
| | **All** | **69** |
| ***P. vivax*** | PRJEB15709 | 5 |
| | PRJNA260605 | 19 |
| | PRJNA279199 | 24 |
| | PRJNA337969 | 55 |
| | PRJNA376620 | 19 |
| | PRJNA378759 | 4 |
| | PRJNA422240 | 4 |
| | PRJNA481383 | 52 |

| | | |
|---|---|---|
| | **All** | **182** |
| ***P. yoelli*** | PRJNA322665 | 16 |
| | PRJNA394583 | 10 |
| | **All** | **26** |
| ***P. berghei*** | PRJEB24219 | 52 |
| | PRJNA286027 | 12 |
| | PRJNA319543 | 124 |
| | PRJNA374918 | 9 |
| | PRJNA390648 | 3 |
| | PRJNA391033 | 9 |
| | PRJNA433164 | 32 |
| | **All** | **241** |
| | **TOTAL** | **1682** |

**Table S3. *Plasmodium vivax* SRA libraries.** Libraries positive for MaRNAV-1 are shown in red and those used for phylogenetic analysis (MaRNAV-1-like read counts > 100) are shown in bold. *Plasmodium*-free libraries are in grey. *Homo sapiens* = human blood samples. *Anopheles dirus* = mosquito salivary gland dissection samples. *Homo sapiens (ex vivo)*: Micropatterned cellular co-cultures. ND: Not determined.

| BioProject | SRA ID | *P. vivax* isolate | Location | Host |
|---|---|---|---|---|
| **PRJNA481383** [(Roth et al. 2018)] | SRR7554416 | *P. vivax* Thai_1 | Thailand | *A. dirus* |
| | SRR7554417 | *P. vivax* Thai_1 | Thailand | *A. dirus* |
| | SRR7554418 | *P. vivax* Thai_1 | Thailand | *A. dirus* |
| | SRR7554419 | *P. vivax* Thai_1 | Thailand | *A. dirus* |
| | SRR7554420 | *P. vivax* Thai_1 | Thailand | *A. dirus* |
| | SRR7554421 | *P. vivax* Thai_1 | Thailand | *A. dirus* |
| | SRR7554422 | *P. vivax* Thai_1 | Thailand | *A. dirus* |
| | SRR7554423 | *P. vivax* Thai_1 | Thailand | *A. dirus* |
| | SRR7554424 | *P. vivax* Thai_1 | Thailand | *A. dirus* |
| | SRR7554425 | *P. vivax* Thai_1 | Thailand | *A. dirus* |
| | SRR7554426 | *P. vivax* Thai_1 | Thailand | *A. dirus* |
| | SRR7554427 | *P. vivax* Thai_1 | Thailand | *A. dirus* |
| | SRR7554428 | *P. vivax* Thai_2 | Thailand | *A. dirus* |
| | SRR7554429 | *P. vivax* Thai_2 | Thailand | *A. dirus* |
| | SRR7554430 | *P. vivax* Thai_2 | Thailand | *A. dirus* |
| | **SRR7554431** | ***P. vivax* Thai_2** | **Thailand** | ***A. dirus*** |
| | SRR7554432 | *P. vivax* Thai_2 | Thailand | *A. dirus* |
| | **SRR7554433** | ***P. vivax* Thai_2** | **Thailand** | ***A. dirus*** |
| | SRR7554434 | *P. vivax* Thai_2 | Thailand | *A. dirus* |
| | **SRR7554435** | ***P. vivax* Thai_2** | **Thailand** | ***A. dirus*** |
| | **SRR7554436** | ***P. vivax* Thai_2** | **Thailand** | ***A. dirus*** |
| | **SRR7554437** | ***P. vivax* Thai_2** | **Thailand** | ***A. dirus*** |
| | **SRR7554438** | ***P. vivax* Thai_2** | **Thailand** | ***A. dirus*** |
| | **SRR7554439** | ***P. vivax* Thai_2** | **Thailand** | ***A. dirus*** |
| | SRR7554440 | *P. vivax* Thai_3 | Thailand | *A. dirus* |
| | **SRR7554441** | ***P. vivax* Thai_3** | **Thailand** | ***A. dirus*** |
| | SRR7554442 | *P. vivax* Thai_3 | Thailand | *A. dirus* |
| | **SRR7554443** | ***P. vivax* Thai_3** | **Thailand** | ***A. dirus*** |
| | SRR7554444 | *P. vivax* Thai_3 | Thailand | *A. dirus* |
| | **SRR7554445** | ***P. vivax* Thai_3** | **Thailand** | ***A. dirus*** |
| | SRR7554446 | *P. vivax* Thai_3 | Thailand | *A. dirus* |
| | **SRR7554447** | ***P. vivax* Thai_3** | **Thailand** | ***A. dirus*** |

| | | | | |
|---|---|---|---|---|
| | SRR7554448 | *P. vivax* Thai_3 | Thailand | *A. dirus* |
| | **SRR7554449** | ***P. vivax*** **Thai_3** | **Thailand** | ***A. dirus*** |
| | SRR7554450 | *P. vivax* Thai_3 | Thailand | *A. dirus* |
| | **SRR7554451** | ***P. vivax*** **Thai_3** | **Thailand** | ***A. dirus*** |
| | SRR7554452 | *P. vivax* Thai_4 | Thailand | *A. dirus* |
| | SRR7554453 | *P. vivax* Thai_4 | Thailand | *A. dirus* |
| | SRR7554454 | *P. vivax* Thai_4 | Thailand | *A. dirus* |
| | SRR7554455 | *P. vivax* Thai_4 | Thailand | *A. dirus* |
| | SRR7554456 | *P. vivax* Thai_4 | Thailand | *A. dirus* |
| | SRR7554457 | *P. vivax* Thai_4 | Thailand | *A. dirus* |
| | SRR7554458 | *P. vivax* Thai_4 | Thailand | *A. dirus* |
| | SRR7554459 | *P. vivax* Thai_4 | Thailand | *A. dirus* |
| | **SRR7554460** | ***P. vivax*** **Thai_5** | **Thailand** | ***A. dirus*** |
| | **SRR7554461** | ***P.vivax*** **Thai_5** | **Thailand** | ***A. dirus*** |
| | **SRR7554462** | ***P.vivax*** **Thai_5** | **Thailand** | ***A. dirus*** |
| | **SRR7554463** | ***P.vivax*** **Thai_5** | **Thailand** | ***A. dirus*** |
| | *SRR7554464* | *Uninfected* | *ND* | *A. dirus* |
| | *SRR7554465* | *Uninfected* | *ND* | *A. dirus* |
| | *SRR7554466* | *Uninfected* | *ND* | *A. dirus* |
| | *SRR7554467* | *Uninfected* | *ND* | *A. dirus* |
| **PRJNA422240** (Gural et al. 2018) | SRR6371894 | *P. vivax* (isolate VK210) #2 | Thailand | *Homo sapiens (ex vivo)* |
| | SRR6371895 | *P. vivax* (isolate VK210) #1 | Thailand | *Homo sapiens (ex vivo)* |
| | SRR6371896 | *P. vivax* (isolate VK210) #2 | Thailand | *Homo sapiens (ex vivo)* |
| | SRR6371897 | *P. vivax* (isolate VK210) #1 | Thailand | *Homo sapiens (ex vivo)* |
| **PRJNA378759** (Kim et al. 2017) | **SRR5646668** | **Sp_1** | **Colombia** | ***Anopheles albimanus*** |
| | **SRR5646669** | **V_DJK_16** | **Cambodia** | ***Homo sapiens*** |
| | SRR5646670 | V_DJK_10 | Cambodia | *Homo sapiens* |
| | SRR5646671 | V_DJK_8 | Cambodia | *Homo sapiens* |
| **PRJNA376620** (Jex et al. ) | SRR5298172 | PvSPZ-Thai9 | Thailand | *A. dirus* |
| | SRR5298173 | PvSPZ-Thai9 | Thailand | *A. dirus* |
| | **SRR5298174** | **PvSPZ-Thai8** | **Thailand** | ***A. dirus*** |

| | | | | |
|---|---|---|---|---|
| | **SRR5298175** | **PvSPZ-Thai8** | **Thailand** | ***A. dirus*** |
| | **SRR5298176** | **PvSPZ-Thai7** | **Thailand** | ***A. dirus*** |
| | **SRR5298177** | **PvSPZ-Thai7** | **Thailand** | ***A. dirus*** |
| | SRR5298178 | PvSPZ-Thai5 | Thailand | *A. dirus* |
| | SRR5298179 | PvSPZ-Thai5 | Thailand | *A. dirus* |
| | **SRR5298180** | **PvSPZ-Thai4** | **Thailand** | ***A. dirus*** |
| | **SRR5298181** | **PvSPZ-Thai4** | **Thailand** | ***A. dirus*** |
| | **SRR5298182** | **PvSPZ-Thai3** | **Thailand** | ***A. dirus*** |
| | **SRR5298183** | **PvSPZ-Thai3** | **Thailand** | ***A. dirus*** |
| | **SRR5298184** | **PvSPZ-Thai2** | **Thailand** | ***A. dirus*** |
| | **SRR5298185** | **PvSPZ-Thai2** | **Thailand** | ***A. dirus*** |
| | SRR5298186 | *P. falciparum (3D7)* | ND | *A. dirus* |
| | SRR5298187 | *P. falciparum (3D7)* | ND | *A. dirus* |
| | **SRR5298188** | **PvSPZ-Thai6** | **Thailand** | ***A. dirus*** |
| | **SRR5298189** | **PvSPZ-Thai6** | **Thailand** | ***A. dirus*** |
| | SRR5298190 | PvSPZ-Thai1 | Thailand | *A. dirus* |
| **PRJNA337969** **(Rojas-Pena et al. )** *P. vivax challenge study* | *SRR4005681* | *Uninfected #10* | - | *Homo sapiens* |
| | *SRR4005682* | *Uninfected #19* | - | *Homo sapiens* |
| | *SRR4005683* | *Uninfected #5* | - | *Homo sapiens* |
| | *SRR4005684* | *Uninfected #7* | - | *Homo sapiens* |
| | *SRR4005685* | *Uninfected #3* | - | *Homo sapiens* |
| | *SRR4005686* | *Uninfected #16* | - | *Homo sapiens* |
| | *SRR4005687* | *Uninfected #1* | - | *Homo sapiens* |
| | *SRR4005688* | *Uninfected #9* | - | *Homo sapiens* |
| | *SRR4005689* | *Uninfected #11* | - | *Homo sapiens* |
| | *SRR4005690* | *Uninfected #18* | - | *Homo sapiens* |
| | *SRR4005691* | *Uninfected #12* | - | *Homo sapiens* |
| | *SRR4005692* | *Uninfected #17* | - | *Homo sapiens* |
| | *SRR4005693* | *Uninfected #13* | - | *Homo sapiens* |
| | SRR4005694 | *P. vivax* - ND #4 | Colombia (Cali) | Homo sapiens |
| | *SRR4005695* | *Uninfected #14* | - | *Homo sapiens* |
| | *SRR4005696* | *Uninfected #6* | - | *Homo sapiens* |
| | *SRR4005697* | *Uninfected #2* | - | *Homo sapiens* |

| | | | |
|---|---|---|---|
| *SRR4005698* | *Uninfected #8* | - | *Homo sapiens* |
| *SRR4005699* | *Uninfected #15* | - | *Homo sapiens* |
| *SRR4005700* | *Uninfected #10* | - | *Homo sapiens* |
| SRR4005701 | *P. vivax* - ND #19 | Colombia (Cali) | *Homo sapiens* |
| *SRR4005702* | *Uninfected #3* | - | *Homo sapiens* |
| *SRR4005703* | *Uninfected #16* | - | *Homo sapiens* |
| *SRR4005704* | *Uninfected #1* | - | *Homo sapiens* |
| *SRR4005705* | *Uninfected #9* | - | *Homo sapiens* |
| SRR4005706 | *P. vivax* - ND #11 | Colombia (Cali) | *Homo sapiens* |
| SRR4005707 | *P. vivax* - ND #12 | Colombia (Cali) | *Homo sapiens* |
| SRR4005708 | *P. vivax* - ND #17 | Colombia (Cali) | *Homo sapiens* |
| SRR4005709 | *P. vivax* - ND #13 | Colombia (Cali) | *Homo sapiens* |
| SRR4005710 | *P. vivax* - ND #4 | Colombia (Cali) | *Homo sapiens* |
| SRR4005711 | *P. vivax* - ND #14 | Colombia (Cali) | *Homo sapiens* |
| *SRR4005712* | *Uninfected #6* | - | *Homo sapiens* |
| *SRR4005713* | *Uninfected #2* | - | *Homo sapiens* |
| *SRR4005714* | *Uninfected #8* | - | *Homo sapiens* |
| *SRR4005715* | *Uninfected #15* | - | *Homo sapiens* |
| *SRR4005716* | *Uninfected #20* | - | *Homo sapiens* |
| *SRR4005717* | *Uninfected #10* | - | *Homo sapiens* |
| *SRR4005718* | *Uninfected #5* | - | *Homo sapiens* |
| *SRR4005719* | *Uninfected #7* | - | *Homo sapiens* |
| *SRR4005720* | *Uninfected #3* | - | *Homo sapiens* |
| *SRR4005721* | *Uninfected #16* | - | *Homo sapiens* |
| *SRR4005722* | *Uninfected #1* | - | *Homo sapiens* |
| *SRR4005723* | *Uninfected #9* | - | *Homo sapiens* |
| *SRR4005724* | *Uninfected #11* | - | *Homo sapiens* |
| *SRR4005725* | *Uninfected #18* | - | *Homo sapiens* |
| *SRR4005726* | *Uninfected #12* | - | *Homo sapiens* |
| *SRR4005727* | *Uninfected #17* | - | *Homo sapiens* |
| *SRR4005728* | *Uninfected #13* | - | *Homo sapiens* |
| SRR4005729 | *P. vivax* - ND #4 | Colombia (Cali) | *Homo sapiens* |
| *SRR4005730* | *Uninfected #14* | - | *Homo sapiens* |

| | | | | |
|---|---|---|---|---|
| | *SRR4005731* | *Uninfected #6* | - | *Homo sapiens* |
| | *SRR4005732* | *Uninfected #2* | - | *Homo sapiens* |
| | *SRR4005733* | *Uninfected #8* | - | *Homo sapiens* |
| | *SRR4005734* | *Uninfected #15* | - | *Homo sapiens* |
| | *SRR4005735* | *Uninfected #20* | - | *Homo sapiens* |
| **PRJNA279199** (Rojas-Peña et al. 2015) *P. vivax challenge study* | SRR1925781 | *P. vivax* - ND #1 | Colombia (Buenaventura) | *Homo sapiens* |
| | *SRR1925782* | *Uninfected #2* | - | *Homo sapiens* |
| | SRR1925783 | *P. vivax* - ND #3 | Colombia (Buenaventura) | *Homo sapiens* |
| | *SRR1925784* | *Uninfected #4* | - | *Homo sapiens* |
| | SRR1925785 | *P. vivax* - ND #5 | Colombia (Buenaventura) | *Homo sapiens* |
| | *SRR1925786* | *Uninfected #1* | - | *Homo sapiens* |
| | SRR1925787 | *P. vivax* - ND #6 | Colombia (Buenaventura) | *Homo sapiens* |
| | SRR1925788 | *P. vivax* - ND #7 | Colombia (Buenaventura) | *Homo sapiens* |
| | *SRR1925789* | *Uninfected #9* | - | *Homo sapiens* |
| | SRR1925790 | *P. vivax* - ND #9 | Colombia (Buenaventura) | *Homo sapiens* |
| | SRR1925791 | *P. vivax* - ND #10 | Colombia (Buenaventura) | *Homo sapiens* |
| | *SRR1925792* | *Uninfected #3* | - | *Homo sapiens* |
| | *SRR1925793* | *Uninfected #11* | - | *Homo sapiens* |
| | *SRR1925794* | *Uninfected #6* | - | *Homo sapiens* |
| | SRR1925795 | *P. vivax* - ND #4 | Colombia (Buenaventura) | *Homo sapiens* |
| | *SRR1925796* | *Uninfected #12* | - | *Homo sapiens* |
| | SRR1925797 | *P. vivax* - ND #12 | Colombia (Buenaventura) | *Homo sapiens* |
| | SRR1925798 | *P. vivax* - ND #2 | Colombia (Buenaventura) | *Homo sapiens* |
| | SRR1925799 | *P. vivax* - ND #8 | Colombia (Buenaventura) | *Homo sapiens* |
| | *SRR1925800* | *Uninfected #9* | - | *Homo sapiens* |
| | *SRR1925801* | *Uninfected #5* | - | *Homo sapiens* |
| | *SRR1925802* | *Uninfected #7* | - | *Homo sapiens* |
| | SRR1925803 | *P. vivax* - ND #11 | Colombia (Buenaventura) | *Homo sapiens* |
| | *SRR1925804* | *Uninfected #10* | - | *Homo sapiens* |
| **PRJNA260605** (Zhu et al. 2016) | SRR1571697 | *P. vivax* - SMRU #1 | Thailand | *Ex vivo* culture |
| | SRR1571698 | *P. vivax* - SMRU #1 | Thailand | *Ex vivo* culture |
| | SRR1571699 | *P. vivax* - SMRU #1 | Thailand | *Ex vivo* culture |
| | SRR1571700 | *P. vivax* - SMRU #1 | Thailand | *Ex vivo* culture |

| | | | | |
|---|---|---|---|---|
| | SRR1571701 | *P. vivax* - SMRU #1 | Thailand | *Ex vivo* culture |
| | SRR1571702 | *P. vivax* - SMRU #1 | Thailand | *Ex vivo* culture |
| | SRR1571703 | *P. vivax* - SMRU #1 | Thailand | *Ex vivo* culture |
| | SRR1571704 | *P. vivax* - SMRU #2 | Thailand | *Ex vivo* culture |
| | SRR1571705 | *P. vivax* - SMRU #2 | Thailand | *Ex vivo* culture |
| | SRR1571706 | *P. vivax* - SMRU #2 | Thailand | *Ex vivo* culture |
| | SRR1571707 | *P. vivax* - SMRU #2 | Thailand | *Ex vivo* culture |
| | SRR1571708 | *P. vivax* - SMRU #2 | Thailand | *Ex vivo* culture |
| | SRR1571709 | *P. vivax* - SMRU #2 | Thailand | *Ex vivo* culture |
| | SRR1571710 | *P. vivax* - SMRU #2 | Thailand | *Ex vivo* culture |
| | SRR1571711 | *P. vivax* - SMRU #2 | Thailand | *Ex vivo* culture |
| | SRR1571712 | *P. vivax* - Mixed | Thailand | *Ex vivo* culture |
| | SRR1571713 | *P. vivax* - Mixed | Thailand | *Ex vivo* culture |
| | SRR1571714 | P. *vivax* - Mixed | Thailand | *Ex vivo* culture |
| | SRR1571715 | *P. vivax* - Mixed | Thailand | *Ex vivo* culture |
| **PRJEB15709** | **ERR1717084** | ***P. vivax*** | **ND** | ***Homo sapiens*** |
| | **ERR1717085** | ***P. vivax*** | **ND** | ***Homo sapiens*** |
| | **ERR1717086** | ***P. vivax*** | **ND** | ***Homo sapiens*** |
| | ERR1717087 | *P. vivax* | ND | *Homo sapiens* |
| | ERR1717452 | *P. vivax* | ND | *Homo sapiens* |

**Table S4. Bird sample analysis summary table.** Presence, absence or uncertainty of *Leucocytozoon* detection from pathology reports are highlighted in green, red and orange, respectively. Positive or negative PCR targeting either the *Leucocytozoon* parasite, the RdRP-like segment and the unknown second segment of MaRNAV-2 are highlighted in green and red, respectively.

| DATE | LOCATION | LEUKOCYTOZOON (Pathology report) | TISSUE | LEUKOCYTOZOON (Mitochondrial ctyB PCR) | MaRNAV-2 (RdRp PCR) | MaRNAV-2 (Segment II PCR) |
|---|---|---|---|---|---|---|
| 2/2/07 | Randwick, NSW | No | Brain | Negative | Negative | Negative |
| | | | Liver | Negative | Positive | Positive |
| | | | Heart | Negative | Negative | Negative |
| 25/11/06 | Centennial Park, NSW | Likely | Brain | Positive | Positive | Positive |
| | | | Kidney | Positive | Positive | Positive |
| 25/11/06 | Kensington, NSW | Likely | Brain | Positive | Positive | Positive |
| | | | Liver | Positive | Positive | Positive |
| 24/8/09 | Seaforth, NSW | No | Brain | Positive | Positive | Positive |
| | | | Heart | Positive | Positive | Positive |
| 30/9/09 | Seaforth, NSW | Yes | Brain | Positive | Positive | Positive |
| | | | Heart | Positive | Positive | Positive |
| | | | Liver | Positive | Positive | Positive |
| 5/1/09 | Sydney, NSW | Yes | Brain | Negative | Positive | Positive |
| | | | Heart | Positive | Positive | Positive |
| | | | Liver | Positive | Positive | Positive |
| 29/10/13 | Nowra, NSW | Yes | Brain | Yes | Positive | Positive |
| | | | Heart | Positive | Positive | Positive |
| 29/10/13 | Nowra, NSW | No | Brain | Negative | Negative | Negative |
| | | | Heart | Negative | Negative | Negative |
| 29/10/13 | Nowra, NSW | Yes | Brain | Yes | Negative | Negative |
| | | | Heart | Yes | Negative | Negative |
| 23/10/13 | Nowra, NSW | No | Brain | Negative | Negative | Negative |
| | | | Heart | Negative | Negative | Negative |
| 28/10/13 | Nowra, NSW | No | Brain | Negative | Negative | Negative |
| | | | Heart | Negative | Negative | Negative |
| 29/10/13 | Mosman, NSW | Yes | Brain | Positive | Positive | Positive |
| | | | Heart | Positive | Positive | Positive |

**Table S5.** Primers used in this study.

| Name | Sequence (5' – 3') | Application |
|---|---|---|
| *P. vivax* _Fw | CGGCTTGGAAGTCCTTGT | *P. vivax and P. falciparum* validation in blood samples. From (Padley et al. 2003) |
| *P. falciparum*_Fw | AACAGACGGGTAGTCATGATTGAG | |
| *Plasmodium*_Rev | GTATCTGATCGTCTTCACTCCC | |
| PkF1140 | GATTCATCTATTAAAAATTTGCTTC | *P. knowlesi* validation in blood samples From (Imwong et al. 2009) |
| PkR1150 | TCTTTTTCTCCGGAGATTAGAACTC | |
| PkF1160 | GATGCCTCCGCGTATCGAC | |
| rPLU3 | TTTTTATAAGGATAACTACGGAAAAGCTGT | *P. knowlesi* validation in blood samples From (Singh et al. 1999) |
| rPLU4 | TACCCGTCATAGCCATGTTAGGCCAATACC | |
| Human_RPS18_Fw | ATGCAGAATCCACGCCAGTA | Human mRNA Detection |
| Human_RPS18_Rev | CCAGACCATTGGCTAGGACC | |
| *Plasmodium*_LDH-P_Fw | GCTTTTCCTTGGGGCATGTT | *Plasmodium* mRNA detection |
| *Plasmodium*_LDH-P_Rev | GGTATGATCGGAGGAGTGATGG | |
| MARNAV-1_Fw_5 | GACTCGTCACCTTGTGAGGC | MARNAV-1 detection (segment I) |
| MARNAV-1_Rev_5 | TGGCATCCACTTCAAGCAGG | |
| MARNAV-1_Fw1 | TTGTCGGTGGAACTCCTTCG | MARNAV-1 full length amplification (segment I) |
| MARNAV-1_Rev4 | ACATCCAAGCAACACACCCT | |
| MARNAV-1_Fw3 | GCCTCACAAGGTGACGAGTC | MaRNAV-1 sequencing (segment I) |
| MARNAV-1_Fw4 | CCTAAGGCGTTCCCTCCTTC | |
| MARNAV-1_Rev1 | CCTGCTTGAAGTGGATGCCA | |
| MARNAV-1_Rev2 | TCCAGTTTTCATCGGCAGGA | |
| MARNAV-1_Rev3 | AAGGCGTCACACCTCAGTAG | |
| MARNAV-1_Fw3 | GCCTCACAAGGTGACGAGTC | |
| Pv_1_unknown_contig Fw1 | GGCGTACTCGTTGCTTTTGT | MARNAV-1 detection, full-length amplification and sequencing (segment II) |
| Pv_1_unknown_contig Rev1 | AATCCTGTCGCGGACACAAT | |
| BW_Narnalike_Fw1 | CTGAAATTGATAARGAYGAAACTCC | Bird samples MARNAV-2 detection (segment I) |
| BW_Narnalike_Rev1 | CGTGGCATCCTTYAAATCTGATG | |
| Haem cytB_AE986-F | AGTGGATGGTGYTTYAGATAYTTAC | Bird samples hematozoa cytB detection From (Pacheco et al. 2018) |
| Haem cytB_AE066-IR | GCTTGGGAGCTGTAATCATAAT | |
| BW.Narna.Novel.3F | TCCATAAATGATGGGAGTAATCGC | Bird samples MARNAV-2 detection (segment II) |
| BW.Narna.Novel.2R | GATCTTGTATATACATAGATCCAATACAG | |

**Table S6.** Quality of RNA extraction and RNA-seq data sets obtained.

| *Plasmodium* species | Sample ID | RNA extraction set | Total RNA quality (RIN) | RNA-Seq depth (read number) |
|---|---|---|---|---|
| ***P. vivax*** | 1 | A | 8.1 | 17,532,644 |
| | 2 | A | 7.4 | |
| | 3 | B | 8 | |
| | 4 | B | 6.2 | |
| | 5 | C | 6.2 | |
| | 6 | C | 6.9 | |
| | 10 | F | 6.3 | |
| ***P. knowlesi*** | 18 | A | 8.2 | 17,082,864 |
| | 19 | A | 8.2 | |
| | 20 | B | 7.4 | |
| | 21 | B | 7.3 | |
| | 22 | C | 6.5 | |
| | 27 | F | 7 | |
| ***P. falciparum*** | 28 | A | 6.9 | 16,666,960 |
| | 31 | C | 7.9 | |
| | 32 | C | 6.6 | |
| | 33 | D | 8.7 | |
| | 35 | E | 7.4 | |
| **Uninfected** | 38 | A | 6.1 | 16,314,125 |
| | 39 | B | 7.1 | |
| | 40 | B | 6.2 | |
| | 42 | C | 7.1 | |
| | 45 | E | 5.9 | |
| | 46 | F | 7 | |

### Table S7. List of databases and software used for rRNA and host read depletion

| Depletion | Reference/accession | Reference link | Software | Software reference |
|---|---|---|---|---|
| **rRNA** | Silva-arc-16s-id95 | (Quast et al. 2013) | SortmeRNA | (Kopylova et al. 2012) |
| | Silva-arc-23s-id98 | | | |
| | Silva-bac-16s-id90 | | | |
| | Silva-bac-23s-id98 | | | |
| | Silva-euk-18s-id95 | | | |
| | Silva-euk-28s-id98 | | | |
| **Short non-coding rRNA** | Rfam-5.8s-database-id98 | (Kalvari et al. 2018; Griffiths-Jones 2005) | | |
| | Rfam-5s-database-id98 | | | |
| **Human** | Refseq \| GCF_000001405.38 | | Bowtie2 | (Langmead and Salzberg 2012) |
| ***P. vivax*** | Refseq \| GCF_000002415.2 | | | |
| ***P. knowlesi*** | Refseq \| GCF_000006355.1 | | | |
| ***P. falciparum*** | Refseq \| GCF_000002765.4 | | | |
